# Mechanism Underlying the Immune Responses of a Sublingual Vaccine for SARS-CoV-2 with RBD Antigen and Adjuvant, Poly(I:C) or AddaS03, in Non-human Primates

**DOI:** 10.1101/2023.05.15.540684

**Authors:** Tetsuro Yamamoto, Fusako Mitsunaga, Kunihiko Wasaki, Atsushi Kotani, Kazuki Tajima, Masanori Tanji, Shin Nakamura

**Author notes:** Correspondence to SN.

## Abstract

A sublingual vaccine formulated with recombinant SARS-CoV-2 spike protein receptor binding domain (RBD) antigen and Poly(I:C)) adjuvant was assessed for its safety in non-human primates. This Poly(I:C)-adjuvanted sublingual vaccine was safe compared to the AddaS03-adjuvanted vaccine in blood tests and plasma CRP. The safety of the vaccine was also confirmed through quantitative reverse transcription PCR of six genes and ELISA of four cytokines associated with inflammation and related reactions. The Poly(I:C)- or AddaS03-adjuvanted sublingual vaccine produced RBD-specific IgA antibodies in nasal washings, saliva, and plasma. SARS-CoV-2 neutralizing antibodies were detected in plasma, suggesting that adjuvanted-sublingual vaccines protect against SARS-CoV-2 infection. “Yin and Yang”-like unique transcriptional regulation was observed through DNA microarray analyses of white blood cell RNAs from both vaccines, suppressing and enhancing immune responses and up- or downregulating genes associated with these immune responses. Poly(I:C) adjuvanted sublingual vaccination induced atypical up- or downregulation of genes related to immune suppression or tolerance; Treg differentiation; and T-cell exhaustion. Therefore, Poly(I:C) adjuvant is safe and favorable for sublingual vaccination and can induce a balanced “Yin/Yang” -like effect on immune responses.

## 1. Introduction

A protein-based vaccine could be routinely used against the severe acute respiratory syndrome coronavirus 2 (SARS-CoV-2) in the pandemic, which is now shifting to an epidemic state. During the coronavirus disease (COVID-19) pandemic, gene-based RNA or DNA vaccines were used to combat this disease; however, these RNA/DNA vaccines had adverse events, including fever, headache, nausea, or chills. [1]. In contrast, protein-based vaccines have fewer side effects and are used against hepatitis and other viral infections [2]. Although time is needed to establish the protein vaccine against SARS-CoV-2, this vaccine is expected to become a mainstay in protecting the world from COVID-19 [3, 4].

An immunity-stimulating adjuvant and protein antigen are indispensable to elicit a protective immune response by this protein-based vaccine. Two types of adjuvants are characterized. The first, MF59 or AS03, is an oil-in-water nano-emulsion vaccine stimulating Th1/Th2 cytokines [5], and the other is a double-stranded (ds)RNA polyinosinic:polycytidylic acid (Poly(I:C)), which is a ligand for Toll-like receptor (TLR) 3 to activate immune and proinflammatory responses [6]. MF59 and AS03 were approved as adjuvants serving as intramuscularly injected vaccines for influenza [7]. However, Poly(I:C) is not approved yet owing to its side effects of fever and proinflammatory cytokine production.

The vaccination mode is also a limiting factor in establishing a protein-based SARS-CoV-2 vaccine. Upper respiratory tract viruses, SARS-CoV-2, and influenza enter the oral or nasal cavity; thus, these mucosae are frequently defined as the first line of defense against pathogens. Mucosal protection generally operates through antibody-mediated and cytotoxic T-cell responses, which can be triggered by mucosal vaccines providing systemic and mucosal responses (locally and at remote mucosal sites). However, sublingual vaccines, like nasal vaccines, induce mucosal immune responses in the upper and lower respiratory tract, stomach, small intestine, and reproductive tract, and a systemic response [8,9]. Another advantage of the sublingual vaccine is that it is safer than the nasal vaccine, which adversely affects the brain, central nervous system, or lungs [10,11]. The sublingual vaccine is also administered needle-free, providing high patient compliance and the possibility of self-administration without medical care personnel. However, vaccine administration via the sublingual route has practical problems, such as a mucin barrier inhibiting vaccine entry into immune cells and a considerable volume of saliva that dilute the vaccine. In a previous study using a non-human primate model, cynomolgus macaques, we improved the mucin inhibition by pre-treatment of the sublingual surface with N-acetyl cysteine (NAC), a mild reducing reagent to disintegrate the mucin layer [12]. We also reduced saliva dilution through anesthesia, a mixture of medetomidine and ketamine, to reduce saliva secretion during vaccine administration [12].

We developed an effective sublingual vaccine formulated with SARS-CoV-2 receptor binding domain (RBD) antigen and Poly(I:C) adjuvant in non-human primates, cynomolgus macaques, in a previous study [12]. In the present study, we aimed to establish the safety of the Poly(I:C) adjuvant for sublingual vaccines by comparing it with the AddaS03 adjuvant, which has the same composition as AS03, in non-human primates. We also investigated the mechanism underlying immune and related responses mediated by the sublingual vaccines.

## 2. Materials and Methods

### 2.1. Reagents and antibodies

NAC, bovine serum albumin, Na-Casein, sodium azide (NaN_3_), and Tween 20 were obtained from Fuji film-Wako (Japan). Phosphate-buffered saline (PBS; Nissui, Japan), polyester swabs (Nipponn Menbou, Japan), filter spin columns (Notgen Biotech), nunc-immune module, F8 Maxisorp (Thermo Fisher Scientific), streptavidin-HRP Conjugate (SA-HRP; Invitrogen), and tetramethyl benzidine (TMB; Sigma-Aldrich) were used. Recombinant SARS-CoV-2 spike protein RBD (Creative Diagnostics), Poly(I:C) HMW vaccine grade ((Poly(I:C); Invitrogen), AddaS03 with a formulation similar to that of the adjuvant system AS03 (Invitrogen), and an ELAST enzyme-linked immunosorbent assay (ELISA) amplification system (PerkinElmer, Inc.) was also used. Biotin-labeled (BT) monkey IgA (Mabtech, Inc.), BT monkey IgA (alpha-chain) (Merck), horseradish peroxidase (HRP)-human IgG (EY Laboratories, Inc), and BT IgE antibodies (Bio-Rad Laboratories, Inc) were used.

RNAiso Plus, PrimeScript™ Reverse Transcriptase, 2680, Recombinant RNase Inhibitor, TB Green® Premix Ex Taq™ (Tli RNaseH Plus), RR420(Takara Bio Inc.), RNeasy MinElute Cleanup Kit (QIAGEN), dNTP Mix and Oligo (dT)15 Primer (Promega), Low Input Quick Amp Labeling Kit, RNA6000 Nano Kit, Agilent Whole Human Genome DNA Microarray 4x44K v2, and Agilent Gene Expression Hybridization Kit (Agilent Technologies) were used in this study.

### 2.2. Animals

Nine cynomolgus macaques (*Macaca fascicularis*; male and female, 12.1–20.6 years old) were used in this study. Following the 3R policy of animal use, the macaque monkeys were reused by subsequential washout for 20 months after termination of some examination(s). These monkeys were negative for B virus, Simian immunodeficiency virus, tuberculosis, *Shigella* spp., *Salmonella* spp., and helminthic parasites.

### 2.3. Vaccination and sampling

Poly(I:C) adjuvant (1 mg/ml) and RBD antigen (2 mg/ml) were stored at -70°C until use. was stored at 4°C until use. Thirteen cynomolgus macaques were divided into three groups: control (nP01∼03), RBD antigen + AddaS03 adjuvant (AddaS03 group; mP04∼08), and RBD antigen + Poly(I:C) adjuvant (Poly(I:C); nP09∼13). The animals in each group were given the following vaccine formula: 0.5 ml of PBS for the control group, 0.5 ml containing 150 µg RBD and 250 µl AddaS03 for the AddaS03 group, and 0.5 ml containing 150 µg RBD and 400 µg Poly(I:C) for the Poly(I:C) group.

The vaccination procedures were conducted under anesthesia with a mixture of medetomidine and ketamine, and atipamezole to wake the monkeys up. Before vaccination, the sublingual surface of the monkeys was pretreated for 5 min using wet cotton dipped in 1% NAC and subsequently washed with saline to disintegrate the mucin layer. After wiping the wet mucin surface with dry cotton, 0.5 ml of each vaccine material, PBS, RBD + AddaS03, or RBD + Poly(I:C) was administered into the sublingual space with a pipette and then allowed to stand for at least 5 min. Sublingual vaccinations were performed three times (0W, 4W, and 8W) at four-week intervals followed by the fourth vaccination after 15 weeks (22W) of the third vaccination as shown in Fig. 1. Blood, saliva, and nasal washings were collected from each monkey under anesthesia as mentioned above. Plasma samples were prepared after centrifuging blood to assay RBD-specific IgA, IgG, or IgE antibodies. Saliva samples were adsorbed to a swab with polystyrene fiber, and nasal washings were recovered using centrifugation in a spin column. Fecal samples collected one day (W0) before and after 7 days of the fourth (W22) vaccination were prepared through extraction, followed by centrifugation. These samples were stored at -40°C until use.

**Figure 1.**
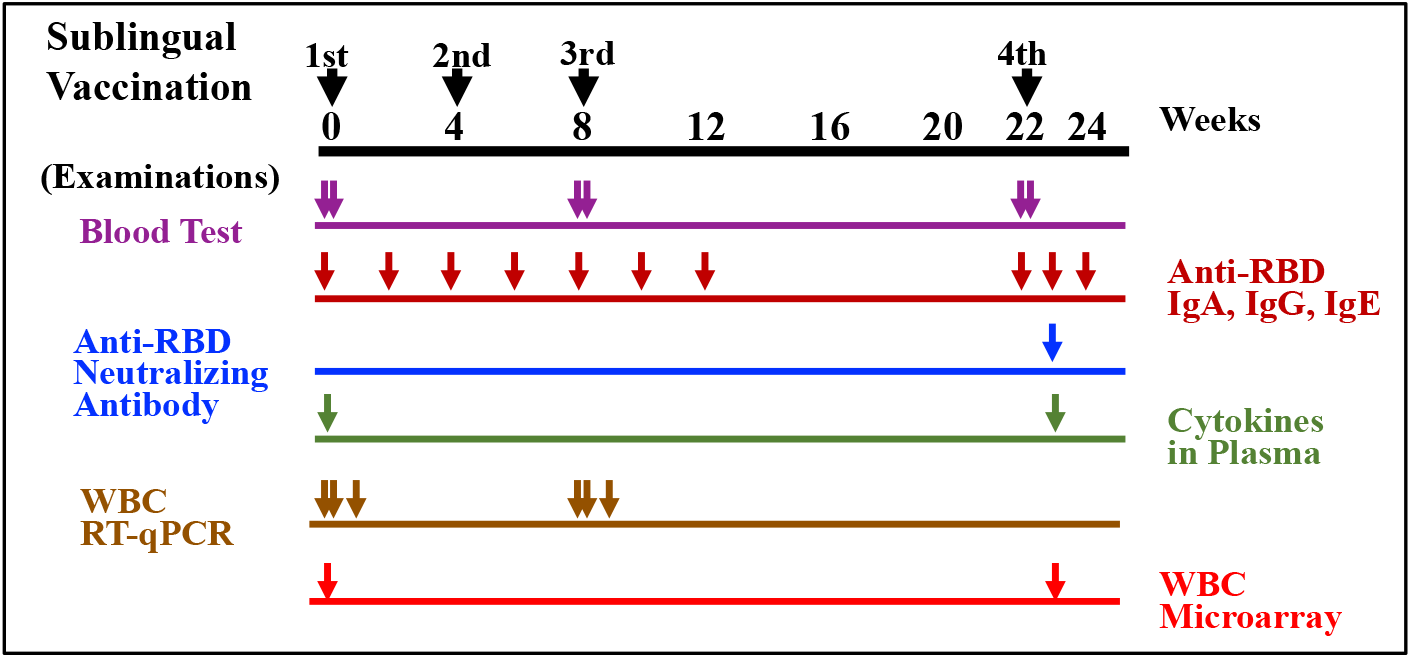
Timepoints of sublingual vaccinations and examinations of blood tests, anti-RBD antibodies, neutralizing antibody, plasma cytokines, WBC RT-qPCR, and WBC microarray. Vaccinations were performed 4 times at 0 (1st), 4 (2nd), 8(3rd), and 22(4th) weeks. Arrows indicate sampling timepoints for each examination.

### 2.4. Blood tests

Blood samples were collected one day before and after the first (W0), third (W8), and fourth (W22) vaccinations, respectively. Fresh whole blood samples were used for the complete blood count of eight items: red blood cells (RBC), white blood cells (WBC), hemoglobin, hematocrit, mean cell volume, mean corpuscular hemoglobin, mean corpuscular hemoglobin concentration, and platelets. Plasma samples were used for biochemical blood tests for total protein, albumin, albumin/globulin ratio, total bilirubin, aspartate transaminase /GOT, alanine transaminase/GPT, alkaline phosphatase, gamma-glutamyl transpeptidase, urea nitrogen-BUN, creatinine, total cholesterol, neutral fats, and C-reactive protein (CRP).

### 2.5. ELISA for RBD-specific antibodies

To detect RBD-specific IgA, IgG, or IgE antibodies, nunc-immune module plates were coated with 100 µl of 5 µg/ml RBD in PBS and incubated at 37°C for 1 h and then at 4°C overnight. After washing with PBS-0.05% Tween 20, 1% Na-Casein in PBS-0.02% NaN_3_ was added to the plates for blocking, followed by incubation at 37°C for 1 h and then stored at 4°C until use. Saliva, nasal washings, and plasma samples were diluted 100 to 500-fold with 1% Na-casein-PBS-0.02% NaN_3._ These diluted samples were used as ELISA samples. To perform an ELISA, after removing the blocking reagent, 50 µl of the diluted ELISA samples and 1M NaCl at a final concentration of 0.5 M were added to the plates to eliminate non-specific reaction.

After incubation at 37°C for 1 h or at 4°C overnight and removing the samples, plates were washed with PBS-0.05% Tween 20. Then, for antibody detection, appropriately diluted BT-monkey IgA, BT-monkey IgA (alpha-chain), HRP-human IgG, or BT-IgE antibodies were added, followed by incubation at 37°C for 1 h. After washing, the plates were amplified using a diluted SA-HRP and an ELAST System mixture comprising biotinyl tyramide. The ELISA sensitivity was enhanced 10 to 30-fold by using this amplification.

After amplification, plates were washed with PBS-0.05% Tween 20, and diluted SA-HRP was added, followed by incubation at 37°C for 1 h. Color development was performed with TMB and terminated by adding H_2_SO_4_. Absorption was measured at 450 and 600 nm using a plate reader (iMark Microplate reader, Bio-Rad Laboratories, Inc).

### 2.6. Neutralizing antibodies

SARS-CoV-2 neutralizing antibody in RBD-specific antibodies was assessed using the SARS-CoV-2 anti-RBD Antibody Profiling Kit (press release by Keio University on September 30, 2020; provided from Medical & Biological Laboratories (MBL)). The neutralizing antibodies in plasma, saliva, or nasal washings were assayed according to the manufacturer’s instructions using samples obtained seven days after the fourth vaccination (W23).

### 2.7. Cytokine assays

Plasma cytokine levels were measured using plasma samples collected one day before and after the first (W0), third (W8), and fourth (W22) vaccinations. Four cytokines, IL-12, IFN-alpha, IFN-gamma, and Il-17, were assayed using ELISA using the Monkey IL12 ELISA Kit (LifeSpan BioSciences, Inc.), Monkey IFN alpha (pan) ELISA Kit (LifeSpan BioSciences Inc), Monkey IFN-gamma ELISA Kit (U-CyTech biosciences), and Monkey IL-17 ELISA Kit (U-CyTech biosciences), respectively.

### 2.8. WBC isolation

Blood samples were collected in heparinized syringes, and WBCs were isolated using erythrocyte lysis buffer, as described by Hoffman et al. [13].

### 2.9. RNA Isolation

Total RNA was extracted from WBCs using the acid guanidine thiocyanate-phenolchloroform method and silica membrane column-based purification [14]. Briefly, WBCs were homogenized in RNAiso Plus (Takara Bio Inc.), and the total RNA was isolated. The RNA was treated with DNase (Qiagen, Valencia, CA, USA) in the aqueous phase and purified using the RNeasy MinElute Cleanup Kit (Qiagen) according to the manufacturer’s instructions. Total RNA (0.2–0.4 mL) of mouse whole blood was isolated using NucleoSpin RNA Blood (Macherey-Nagel GmbH & Co.) according to the manufacturer’s instructions with on-column DNA digestion. The quantity and purity of RNA were evaluated at 230, 260, 280, and 320 nm using an Ultrospec 2000 spectrometer (GE Healthcare Bio-Sciences AB, Uppsala, Sweden).

For DNA microarray analyses, RNA integrity number (RIN) was determined using an Agilent 2100 Bioanalyzer (Agilent Technologies Japan Ltd., Tokyo, Japan). Only high-quality RNA samples, which were defined by an A260/A230 of 1.5, A260/A280 of 1.8, and RIN 6.0, were used in the microarray analysis.

### 2.10. Gene expression analyses using quantitative reverse transcription PCR

The mRNA levels of target genes in WBC samples were determined using quantitative reverse transcription PCR (RT-qPCR), as described previously [15]. cDNA was synthesized from the purified RNA using PrimeScript Reverse Transcriptase with RNase Inhibitor (Takara Bio Inc.), dNTP mixture (Promega Corp., Madison, WI, USA.), and Oligo dT primers (Invitrogen, Waltham, MA, USA). Real-time PCR was performed using the Mx3000P QPCR System (Agilent Technologies Inc., Santa Clara, CA, USA) with a SYBR Premix Ex Taq II (Tli RNaseH Plus) Kit (Takara Bio Inc.). Specific primers for monkey IL12a, IL12b, GZMB, type I IFNs (IFN-alpha1 and IFN-beta1), CD69, and the reference gene low-density lipoprotein receptor-related protein 10 (Lrp10) were constructed using Primer3 and Primer-BLAST [15]. A standard curve was generated by serial dilution of a known amount of glyceraldehyde 3-phosphate dehydrogenase amplicon to calculate the cDNA copy number of the genes. The PCR conditions were 95°C for 15 s, followed by 35 cycles of 95°C for 10 s and 63°C for 30 s, with a dissociation curve. The quantity of target gene mRNA was expressed as the ratio against that of a suitable reference gene, Lrp10 [16].

### 2.11. DNA microarray analyses

DNA microarray analyses were performed for the three groups, i.e., the control, Poly(I:C), and AddaS03. After cDNA synthesis, Cy3-labeled cRNA was synthesized and purified using the Low Input Quick Amp Labeling Kit (Agilent) according to the manufacturer’s instructions. Notably, reverse transcription was conducted using a T7 promoter-oligo(dT) primer. Absorbance was measured at 260, 280, 320, and 550 nm, and it was verified that the labeled cRNA had incorporated >6 pmol/mg of Cy3-CTP. Labeled cRNA was fragmented using the Gene Expression Hybridization Kit (Agilent) and applied to Whole Mouse Genome Array Ver2.0 slides (Agilent). After hybridization at 65 °C for 17 h, the slides were washed with Gene Expression Wash Buffers 1 and 2 (Agilent) according to the manufacturer’s instructions. The slides were scanned using a GenePix 4000B scanner (Molecular Devices Japan K.K., Tokyo, Japan). Scanned images were digitalized and normalized using GebePix Pro software (Molecular Devices Japan K.K., Tokyo, Japan).

### 2.12. Bioinformatic analysis of microarray data

By comparing the control group vs. the Poly(I:C) and AddaS03 groups, or control group vs. the Poly(I:C) or AddaS03 group, a list of genes that were >2-fold upregulated or <0.5-fold downregulated in the Poly(I:C) and AddaS03 groups or the Poly(I:C) or AddaS03 group were generated. We referred to these genes as vaccine-associated genes. The top 50 vaccine-associated genes were annotated, and references were searched using NCBI databases, Google, and related information. The immune-related functions of annotated genes were deduced regarding the possible mechanisms of action of the sublingual vaccine with a Poly(I:C) or AddaS03 adjuvant.

## 3. Results

### 3.1 Safety

Vaccine safety was assessed using blood tests and blood counts, and biochemical tests. Blood or plasma samples from three groups (control, Poly(I:C), and AddaS03) was subjected to the blood tests one day pre- and post-vaccination at three time points: first (W0), third (W8), and fourth (W22) vaccinations (Fig. 1). Between the control and two vaccinated groups, the level of the tests were almost at the same range at any time point (data not shown). The plasma CRP level were also determined for the control and two vaccinated groups, and the level of the three groups increased one-day post-vaccination (data not shown), suggesting CRP increase by experimental stress but not by vaccination. These indicate that sublingual vaccination with Poly(I:C) adjuvant is as safe as with the AddaS03 adjuvant.

### 3.2. RBD-specific antibodies

Figure 2 shows the RBD-specific antibody titer by sublingual vaccination, anti-RBD IgA in saliva, nasal washings, anti-RBD IgA, and anti-RBD IgG in plasma. RBD-specific IgA antibody was detected in saliva and nasal washings after the third vaccination in the Poly(I:C) and AddaS03 groups (Figs. 2A and 2B). The production pattern of RBD-specific IgA, IgG antibodies, or both in the plasma differed from that in saliva and nasal washings. In the Poly(I:C) and AddaS03 groups, plasma anti-RBD IgA and IgG antibodies were undetectable by the third vaccination but detectable after the fourth vaccination (Figs. 2C and 2D). Neither anti-RBD IgA and IgG antibodies in fecal samples nor anti-RBD IgE plasma were detected after one week from the fourth vaccination (data not shown).

**Figure 2.**
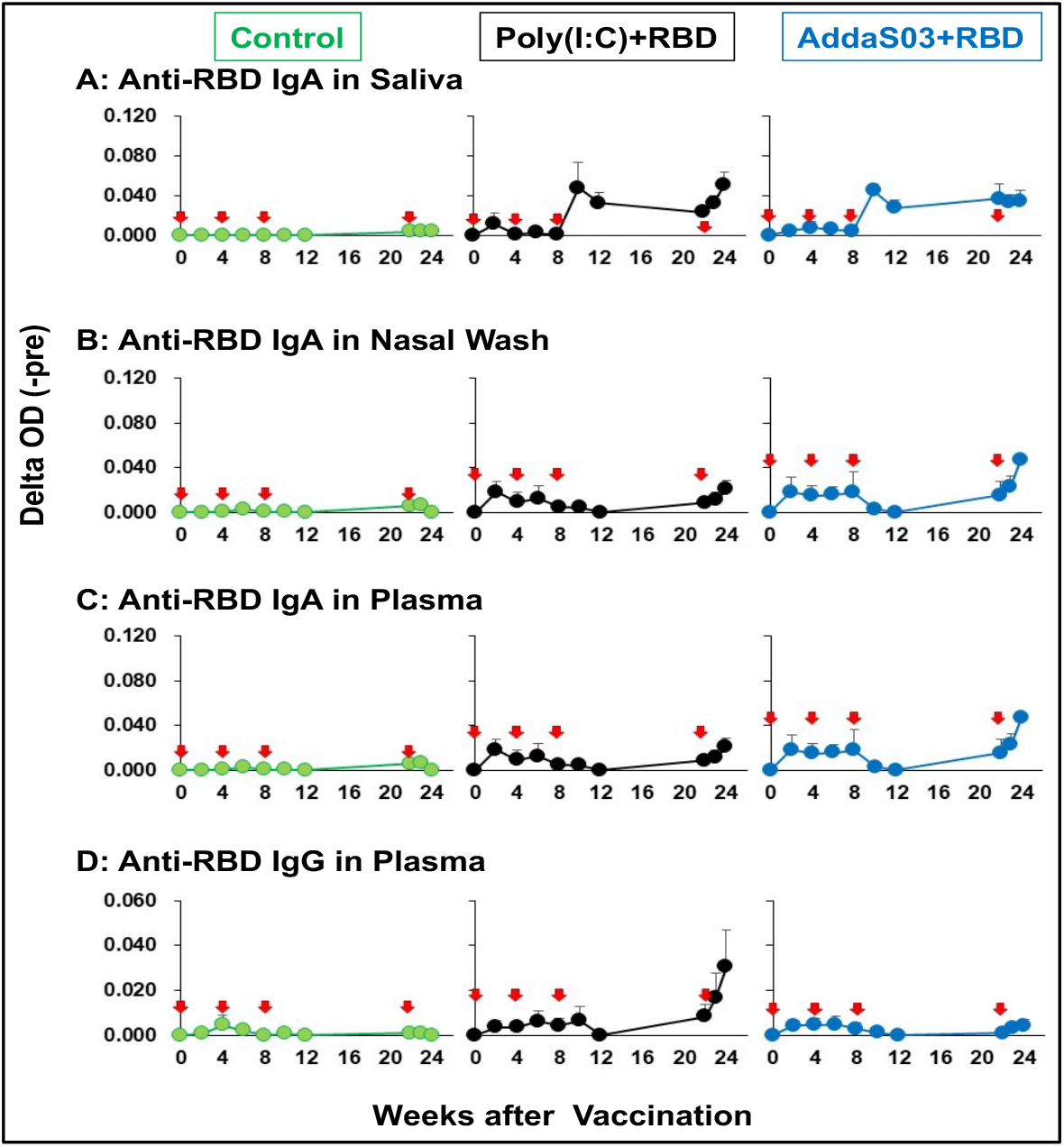
RBD-specific antibodies induced by sublingual vaccination with SARS-CoV-2 RBD + Poly(I:C) or AddaS03 adjuvant. Red arrows indicate vaccination.

These results suggest that a sublingual vaccine with a Poly(I:C) or AddaS03 adjuvant elicits an immune response to produce antigen-specific antibodies in nasal and oral cavities and the blood. These sublingual vaccines appear to act on different mechanisms of the immune system to produce antigen-specific antibodies in local mucosal and systemic immune systems.

### 3.3 Neutralizing antibodies

Plasma samples obtained seven days after the fourth vaccination (W23, Fig. 1) were assayed using the SARS-CoV-2 anti-RBD Antibody Profiling Kit to determine SARS-CoV-2 neutralizing antibody production in the Poly(I:C) and AddaS03 groups. As shown in Fig.3, neutralizing antibodies in plasma were detected semi-dose dependently with anti-RBD IgA antibody titer. Although little neutralizing activity in the saliva or nasal washings was detected (data not shown), these are probably caused by existing of an inhibitor factor, soluble form of ACE2, in saliva or nasal washings as mentioned in Dscussion.

**Figure 3.**
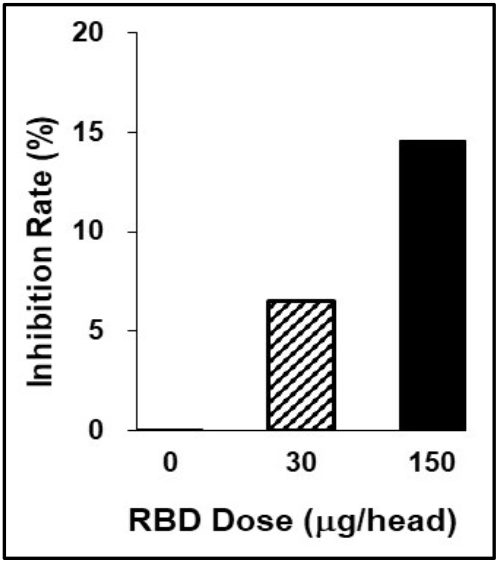
Neutralizing activity of plasma samples form the Poly(I:C) group. Inhibition rate indicates the ability of antibodies to inhibit the interaction between ACE2 and SARS-CoV-2 RBD as described in protocol of the kit used here.

### 3.4. Cytokine production

To assess immune-proinflammatory responses in the Poly(I:C) and AddaS03 groups, we examined four cytokines, IFN-alpha, IFN-gamma, IL-12, and IL-17. These cytokines were assayed using ELISA and plasma samples from pre-vaccination (W0) and 7 days after the fourth vaccination (W23). As shown in Fig. 4, the relative plasma level of IFN-alpha, IFN-gamma, and IL-12 was the same between the Poly(I:C) and AddaS03 groups, in which no increase of these cytokines was observed in comparison with those of control. Notably, a 1.5-fold higher IL-17 level was observed in the Poly(I:C) group, suggesting that the Poly(I:C) adjuvant has a much more potent effect on activating Th17 cells than AddaS03.

**Figure 4.**
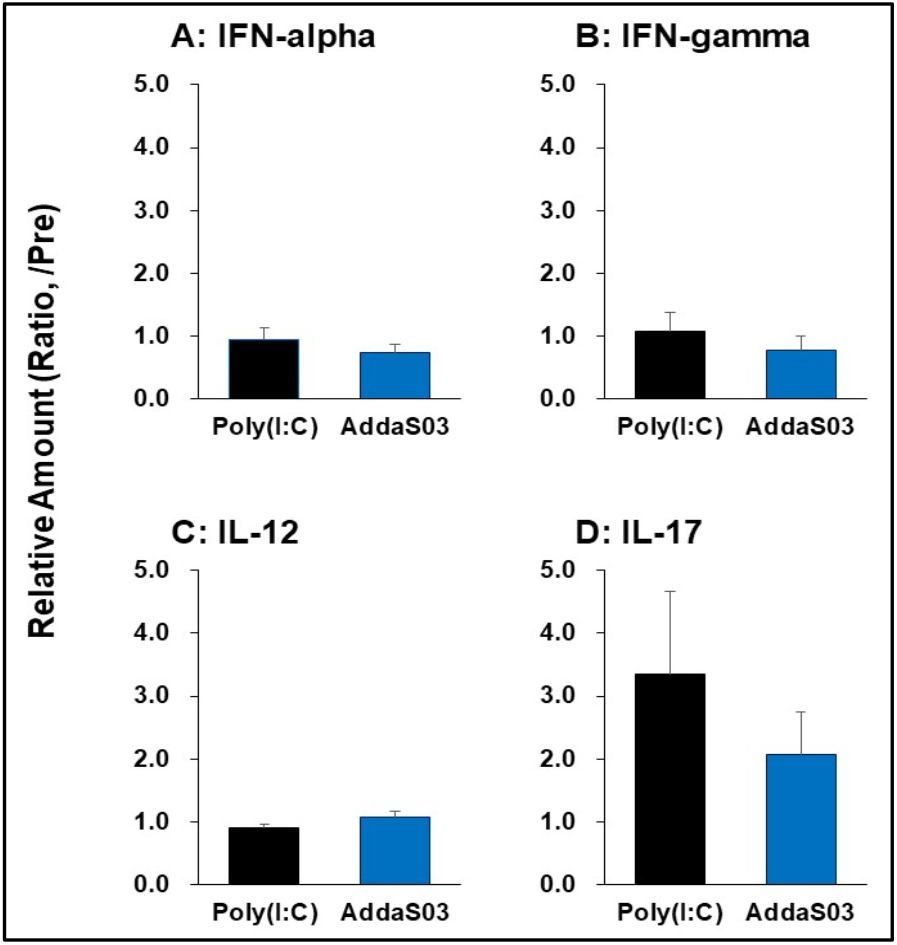
Cytokine production in RBD + Poly(I:C) and RBD + AddaS03 groups. Changes of cytokine amount in plasma after 4th vaccination were expressed as ratio against those before vaccination.

### 3.5. RT-qPCR of immuno-proinflammatory factors

We examined gene expression analyses of six immuno-proinflammatory factors: IL12a; IL12b; GZMB, IFN-alpha1, IFN-beta1, and CD69. The gene expression analyses were performed using RT-qPCR employing primers and RNAs from WBCs of the control, Poly(I:C), and AddaS03 groups. WBCs were collected at three time points (Fig. 1); pre-vaccination (W0/D0 and W8/D56), 1day post-vaccination (W0/D0 and W8/D56), and 1 week after (W1/D7and W9/D63) the first and third vaccinations, respectively. As shown in Fig.5, a similar pattern of relative gene expression of the 6 genes was observed after the first and third vaccinations of the Poly(I:C) and AddaS03 groups. The expression of these genes was upregulated 1 day after vaccination, and then their upregulated expression returned to the level of prevaccination at 7 days after the vaccination. Slightly higher gene expression was observed in IL12A, CZMB, IFN-alpha1, IFN-beta1, and CD69 in the AddaS03 group. These results suggest that in the sublingual vaccination Poly(I:C) adjuvant had a milder effect on immuno-proinflammatory factor than AddaS03, comprising the same components as those of a safe vaccine adjuvant, AS03. Thus, the Poly(I:C) adjuvant is considered safe in sublingual vaccination judging from gene expression in WBCs.

**Figure 5.**
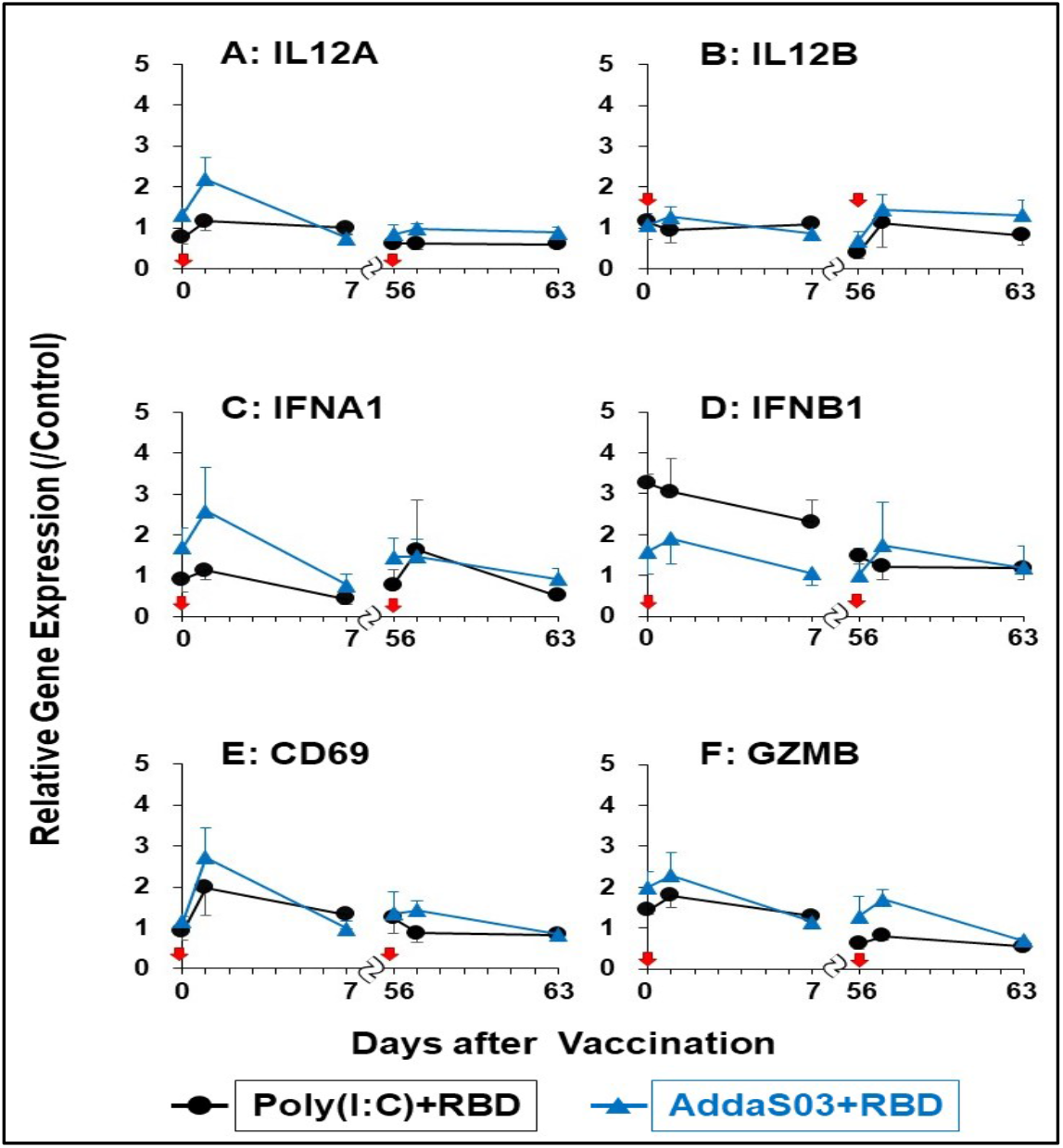
Changes of mRNA level of cytokines and effector molecules in RBD + Poly(I:C) and RBD + AddaS03 groups ed arrows indicate vaccination.

**Figure 6.**
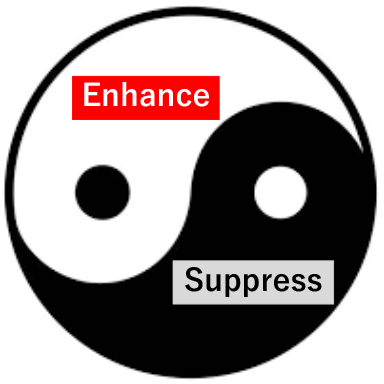
Yin/Suppress and Yang/Enhance-like balance in immune responses elicited by sublingual vaccination using Poly(I:C) adjuvant.

### 3.6. DNA microarray analyses and bioinformatics

We deduced the immune response mechanism from sublingual vaccination with the Poly(I:C) or AddaS03 adjuvant. We also clarified the difference in transcriptional events in the Poly(I:C) or AddaS03 groups. Therefore, we performed DNA microarray analyses using WBC samples at pre-vaccination (W0), and seven days after the fourth vaccination (W23) of the control, Poly(I:C), or AddaS03 groups (Fig. 1). Tables 1 to 3 present genes upregulated more than 2-fold in each group with a comparison of gene expression to the control group. Tables 4 to 6 show genes downregulated less than half in each group, comparing gene expression to the control group.

**Table 1.** Immune related genes upregulated more than 2-fold in both Poly(I:C) and AddaS03 groups.

| Gene symbol | Fold change |  | Product; Description; Function [References] | Expected effect** |
| --- | --- | --- | --- | --- |
|  | Poly(I:C) | AddaS03 |  |  |
| *CCL7\** | 11.6 | 2.2 | C-C motif chemokine ligand 7 (MCP3); Monocyte migration to inflammatory site [17] | ↑ |
| *CCL2\** | 8.0 | 2.3 | C-C motif chemokine ligand 2(MCP1); Monocyte migration to inflammatory site | ↑ |
| *TNFRSF12A\** | 5.9 | 4.2 | TNF receptor superfamily member 12A; Inhibits T-cell proliferation & activation [18] | ↓ |
| *RGS1\** | 4.4 | 3.1 | Regulator of G protein signaling 1; marker of T-cell exhaustion [19, 20] | ↓ |
| *SLA\** | 4.0 | 2.5 | Src like adaptor protein; Inhibits T-cell proliferation and activation [21] | ↓ |
| *ADM* | 3.4 | 2.4 | Adrenomedullin; Angiogenesis and antimicrobial activity |  |
| *CXCR4\** | 3.4 | 2.5 | C-X-C motif chemokine receptor 4; B-cell generation to antibody production [22] | ↑ |
| *PFKFB3\** | 2.9 | 2.3 | 6-Phosphofructo-2-kinase/Fructose-2,6-biphosphatase 3; Glucose metabolism, T-cell activation, & Macrophage survival [23-25] | ↑ |
| *JUN\** | 2.7 | 2.3 | Jun proto-oncogene; AP-1 formation. T-cell activation [26, 27] | ↑ |
| *B3GNT5* | 2.7 | 2.9 | Beta-Gal beta-1,3-N-acetylglucosaminyltransferase 5; |  |
| *PTX3\** | 2.5 | 2.1 | Pentraxin 3; IL-6-induced acute inflammatory response protein [28] | ↑ |
| *FADD\** | 2.4 | 2.3 | Fas associated via death domain; NF-kB activation. T-cell activation [29] | ↑ |
| *ETV6\** | 2.4 | 2.4 | ETS Variant Transcription Factor 6; Stimulation of CD4+T-cell antigen memory [30] | ↑ |
| *KLHL2\** | 2.1 | 2.5 | Kelch like family member 2; Induces JUN gene expression [31] | ↑ |
| *STX11* | 2.1 | 2.2 | Syntaxin 11; Release of lytic granule from NK cells. |  |
| *LRRC8C* | 2.0 | 2.1 | Leucine rich repeat containing 8 VRACsubunit C |  |

Upregulation fold was estimated by relative gene expression calculated from that of pre-vaccination (W0) and post-7 days (W23) of 4th vaccination. \*These genes are mentioned in results and discussion. \*\*Expected effect means that the gene expression change, upregulation or downregulation, is expected to enhance (↑) or suppress (↓) immune and its related responses.

#### 3.6.1. Upregulated genes

Table 1 shows 16 immune-related genes upregulated in the Poly(I:C) and AddaS03 groups. Of these, 12 typical genes were identified; ***CCL7*** and ***CCL2***: a chemoattractant for several leukocytes to mediate immune response, ***TNFRSF12A***: TWWEAK receptor to regulate T-cell activation and proliferation, ***RGS1***: regulates T-cell proliferation and a marker of T cell exhaustion, ***SLA***: inhibits signal transduction through T cell receptors, ***CXCR4*:** crucial action on B-cell generation to antibody production, ***PFKFB*3**: stimulates T cell activation by activating the glycolytic pathway, *JUN*: induces CCL2 and stimulates several immune responses, ***PTX3***: inflammatory factor induced by IL-6, ***FADD***: activates NF-kB, signal transduction, and T-cells, ***ETV6***: memory reading of CD4^+^ memory T cells, and ***KLHL2***: induction of the *Jun* gene and antigen processing/presentation. Table 2 shows 28 genes upregulated in only the Poly(I:C) group. Seven characteristic genes were upregulated; ***ZNF16***: TGF-beta pathway activation, ***EDN1***: TGF-beta pathway activation, ***C15orf48***: simulation of the inflammatory response, ***SLC2A1***: T-cell activation through Glut2 increase, ***PRDM1***: T cell exhaustion and Treg differentiation, ***NR4A3***: T cell exhaustion, and ***EWSR1***: B cell response regulation and antibody production.

**Table 2.**
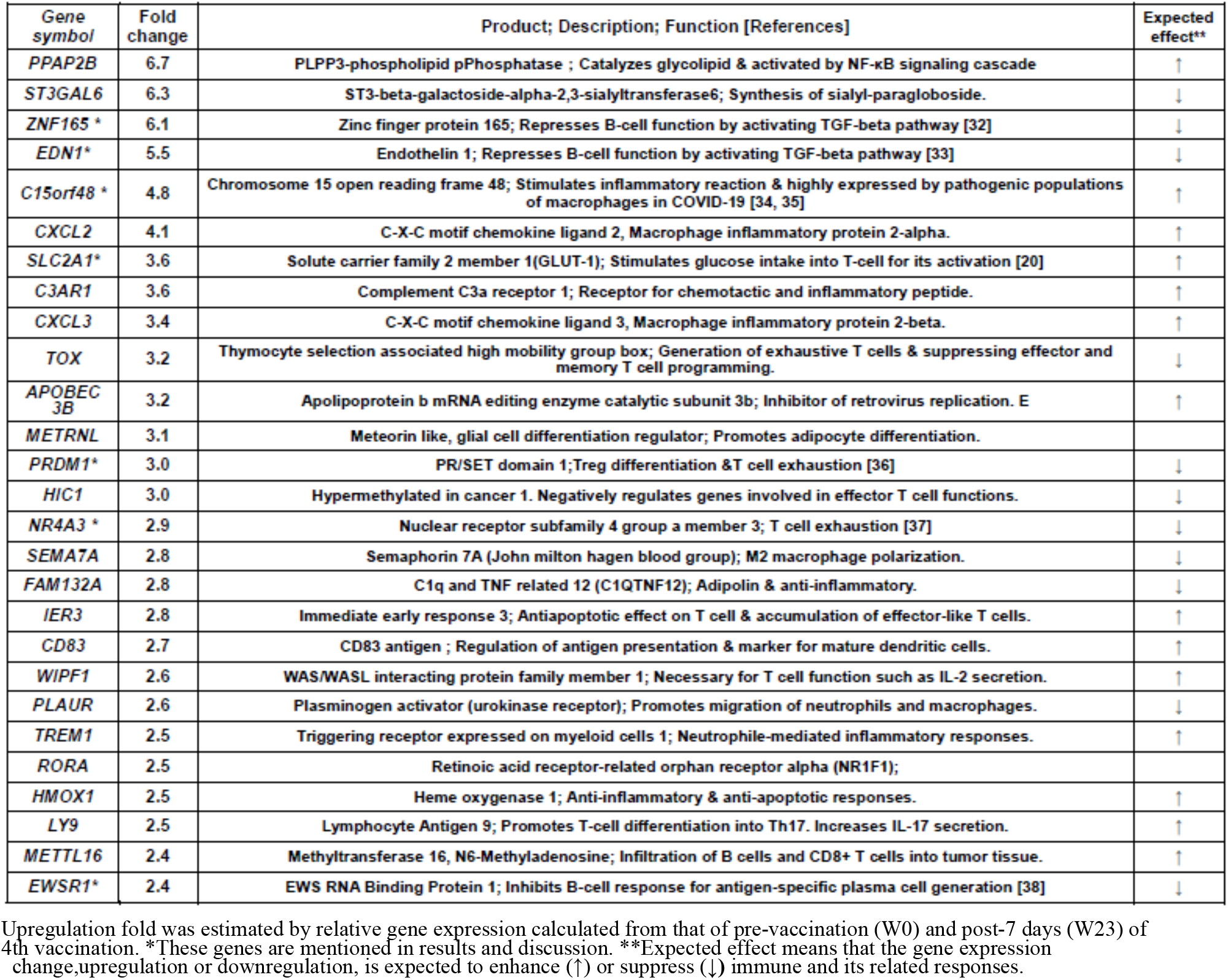
Immune-related genes upregulated more than 2-fold in Poly(I:C) group alone.

Table 3 shows 13 genes upregulated in only the AddaS03 group. Within these, seven marked genes were as follows; ***PDK4***: B-cell differentiation by suppressing CD20 production, ***IFIT1, 2, 3***: anti-virus activity and type I IFN induction, ***GBP1***: anti-bacterial activity and induced by IFN, ***ISG15***: IFN-inducible ubiquitin-like protein and NK cell activation, and ***GAB2***: inhibits T cell receptor-mediated signal transduction.

**Table 3.**
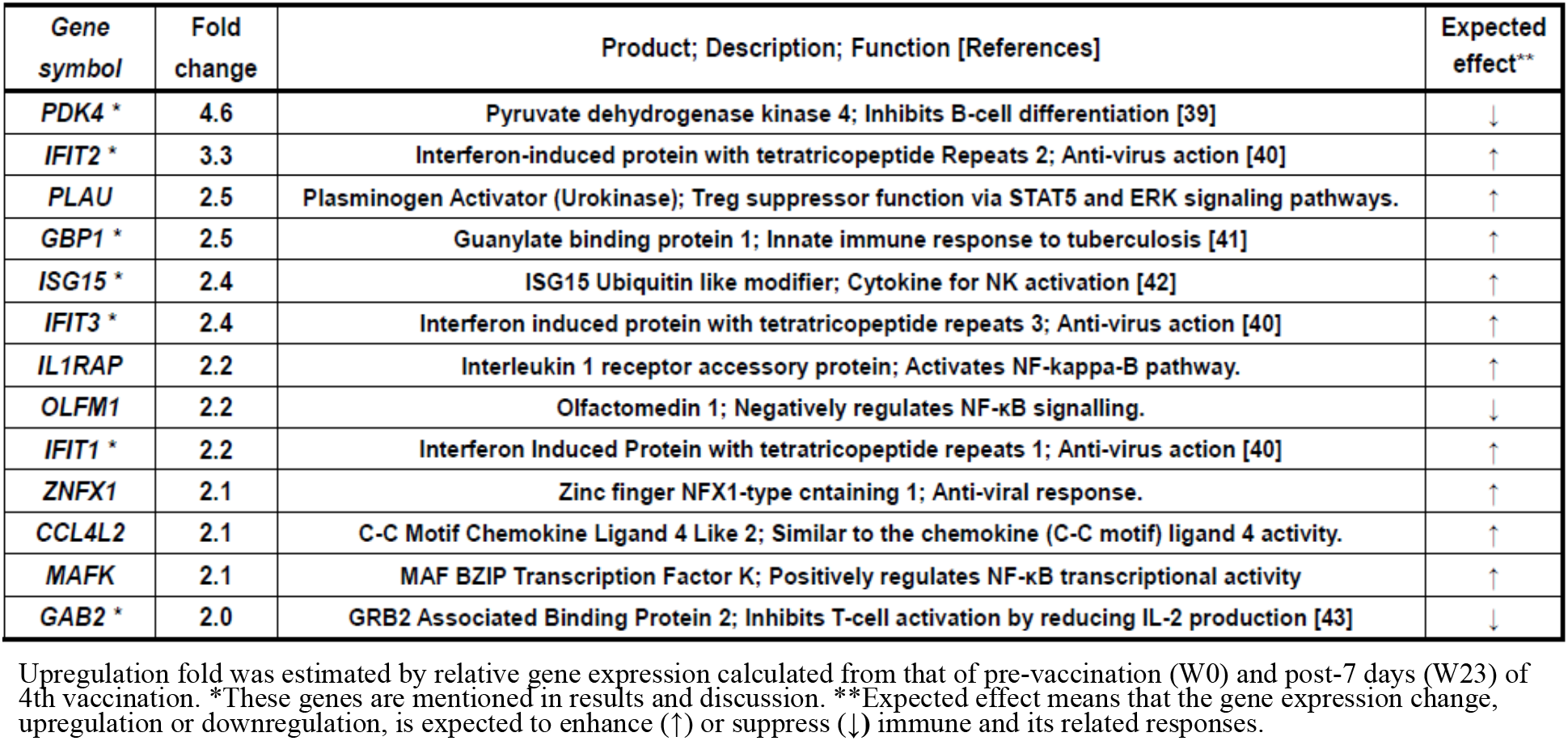
Immune-related genes upregulated more than 2-fold in AddaS03 group alone.

| Gene symbol | Fold change | Product; Description; Function [References] | Expected effect** |
| --- | --- | --- | --- |
| **PDK4 \*** | 4.6 | Pyruvate dehydrogenase kinase 4; Inhibits B-cell differentiation [39] | ↓ |
| **IFIT2 \*** | 3.3 | Interferon-induced protein with tetratricopeptide Repeats 2; Anti-virus action [40] | ↑ |
| **PLAU** | 2.5 | Plasminogen Activator (Urokinase); Treg suppressor function via STAT5 and ERK signaling pathways. | ↑ |
| **GBP1 \*** | 2.5 | Guanylate binding protein 1; Innate immune response to tuberculosis [41] | ↑ |
| **ISG15 \*** | 2.4 | ISG15 Ubiquitin like modifier; Cytokine for NK activation [42] | ↑ |
| **IFIT3 \*** | 2.4 | Interferon induced protein with tetratricopeptide repeats 3; Anti-virus action [40] | ↑ |
| **IL1RAP** | 2.2 | Interleukin 1 receptor accessory protein; Activates NF-kappa-B pathway. | ↑ |
| **OLFM1** | 2.2 | Olfactomedin 1; Negatively regulates NF-κB signalling. | ↓ |
| **IFIT1 \*** | 2.2 | Interferon Induced Protein with tetratricopeptide repeats 1; Anti-virus action [40] | ↑ |
| **ZNFX1** | 2.1 | Zinc finger NFX1-type containing 1; Anti-viral response. | ↑ |
| **CCL4L2** | 2.1 | C-C Motif Chemokine Ligand 4 Like 2; Similar to the chemokine (C-C motif) ligand 4 activity. | ↑ |
| **MAFK** | 2.1 | MAF BZIP Transcription Factor K; Positively regulates NF-κB transcriptional activity | ↑ |
| **GAB2 \*** | 2.0 | GRB2 Associated Binding Protein 2; Inhibits T-cell activation by reducing IL-2 production [43] | ↓ |

Upregulation fold was estimated by relative gene expression calculated from that of pre-vaccination (W0) and post-7 days (W23) of 4th vaccination. \*These genes are mentioned in results and discussion. \*\*Expected effect means that the gene expression change, upregulation or downregulation, is expected to enhance (↑) or suppress (↓) immune and its related responses.

#### 3.6.2. Downregulated genes

As shown in Table 4, seventeen genes were downregulated in the Poly(I:C) and AddaS03 groups. Of these, eight distinctive genes were downregulated; ***PPBP***: CXCL7 chemokine, ***ITGB5***: stimulates M2 macrophage polarization, ***AQP3***: IL-6 and TNF-alpha production in macrophages, ***AQP1***: IL1-beta production in macrophages, ***GP9*** and ***GP1BB***: blood coagulation and neutrophil extracellular trap, ***C6orf25***: CD4^+^ T-cell repression, and ***TESPA1***: T-cell differentiation in the thymus. Table 5 shows 25 genes downregulated in Poly(I:C) groups. Among these genes, eight characteristic genes included; ***TNFAIP6***: macrophages transition to anti-inflammatory phenotype, ***HSPA1B***: stimulation of antigen presentation, ***WIPI*1**: autophagy function, ***IKBIP***: NF-kB pathway inhibition, ***TNFSF1/TRAIL***: apoptosis induction, ***NREP***: TGF-beta production stimulation, ***PGLYRP1***: cytotoxic activity to target cells, and ***IL27RA***: repression of Th2 subset generation. Table 6 lists 18 genes downregulated in Poly(I:C) groups.

**Table 4.**
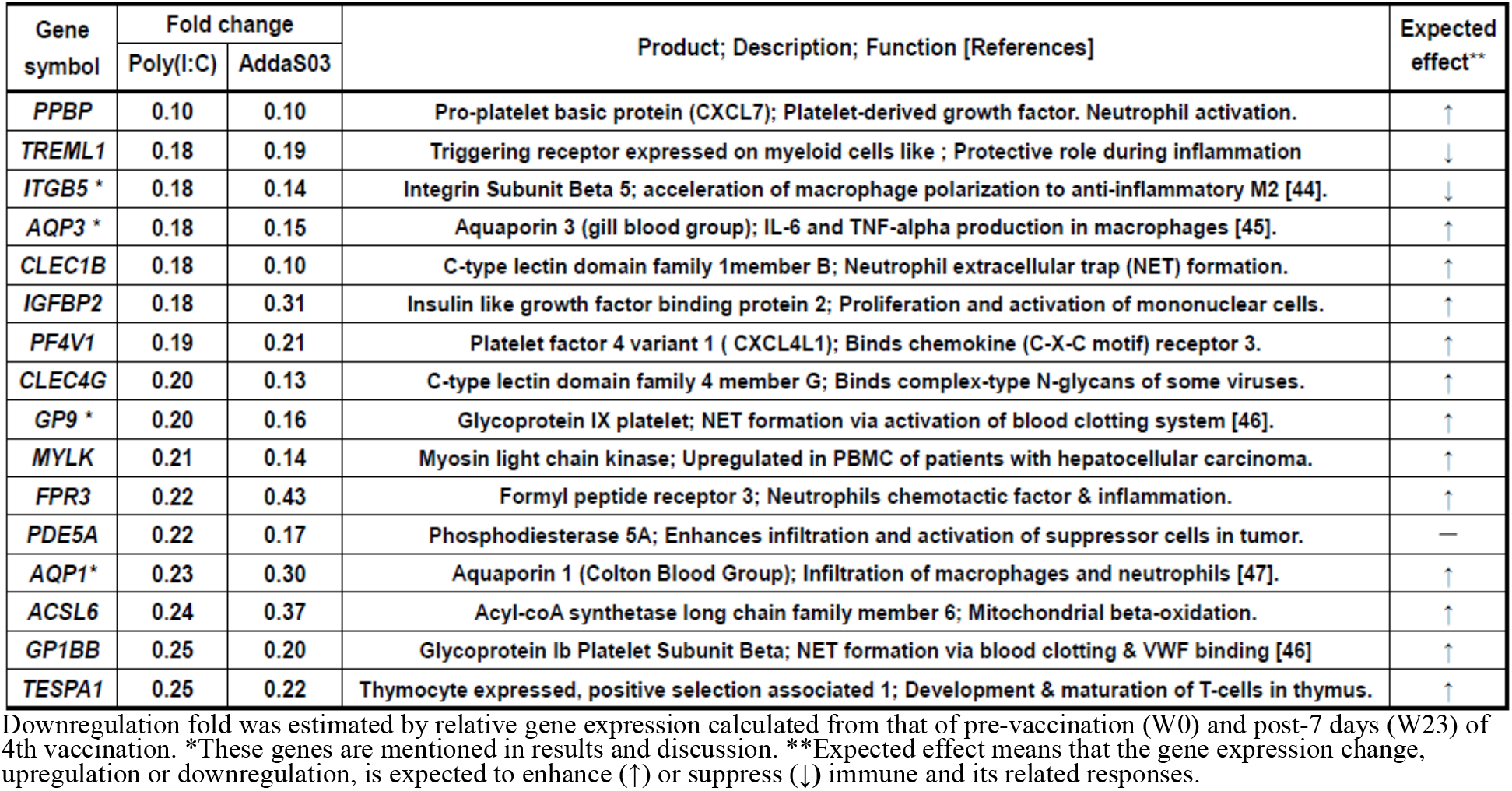
Immune-related genes downregulated less than 0.5-fold both in Poly(I:C) and AddaS03 groups

| Gene symbol | Fold change |  | Product; Description; Function [References] | Expected effect** |
| --- | --- | --- | --- | --- |
|  | Poly(I:C) | AddaS03 |  |  |
| **PPBP** | 0.10 | 0.10 | Pro-platelet basic protein (CXCL7); Platelet-derived growth factor. Neutrophil activation. | ↑ |
| **TREML1** | 0.18 | 0.19 | Triggering receptor expressed on myeloid cells like ; Protective role during inflammation | ↓ |
| **ITGB5 \*** | 0.18 | 0.14 | Integrin Subunit Beta 5; acceleration of macrophage polarization to anti-inflammatory M2 [44]. | ↓ |
| **AQP3 \*** | 0.18 | 0.15 | Aquaporin 3 (gill blood group); IL-6 and TNF-alpha production in macrophages [45]. | ↑ |
| **CLEC1B** | 0.18 | 0.10 | C-type lectin domain family 1member B; Neutrophil extracellular trap (NET) formation. | ↑ |
| **IGFBP2** | 0.18 | 0.31 | Insulin like growth factor binding protein 2; Proliferation and activation of mononuclear cells. | ↑ |
| **PF4V1** | 0.19 | 0.21 | Platelet factor 4 variant 1 ( CXCL4L1); Binds chemokine (C-X-C motif) receptor 3. | ↑ |
| **CLEC4G** | 0.20 | 0.13 | C-type lectin domain family 4 member G; Binds complex-type N-glycans of some viruses. | ↑ |
| **GP9 \*** | 0.20 | 0.16 | Glycoprotein IX platelet; NET formation via activation of blood clotting system [46]. | ↑ |
| **MYLK** | 0.21 | 0.14 | Myosin light chain kinase; Upregulated in PBMC of patients with hepatocellular carcinoma. | ↑ |
| **FPR3** | 0.22 | 0.43 | Formyl peptide receptor 3; Neutrophils chemotactic factor & inflammation. | ↑ |
| **PDE5A** | 0.22 | 0.17 | Phosphodiesterase 5A; Enhances infiltration and activation of suppressor cells in tumor. | — |
| **AQP1\*** | 0.23 | 0.30 | Aquaporin 1 (Colton Blood Group); Infiltration of macrophages and neutrophils [47]. | ↑ |
| **ACSL6** | 0.24 | 0.37 | Acyl-coA synthetase long chain family member 6; Mitochondrial beta-oxidation. | ↑ |
| **GP1BB** | 0.25 | 0.20 | Glycoprotein Ib Platelet Subunit Beta; NET formation via blood clotting & VWF binding [46] | ↑ |
| **TESPA1** | 0.25 | 0.22 | Thymocyte expressed, positive selection associated 1; Development & maturation of T-cells in thymus. | ↑ |

Downregulation fold was estimated by relative gene expression calculated from that of pre-vaccination (W0) and post-7 days (W23) of 4th vaccination. \*These genes are mentioned in results and discussion. \*\*Expected effect means that the gene expression change, upregulation or downregulation, is expected to enhance (↑) or suppress (↓) immune and its related responses.

**Table 5.**
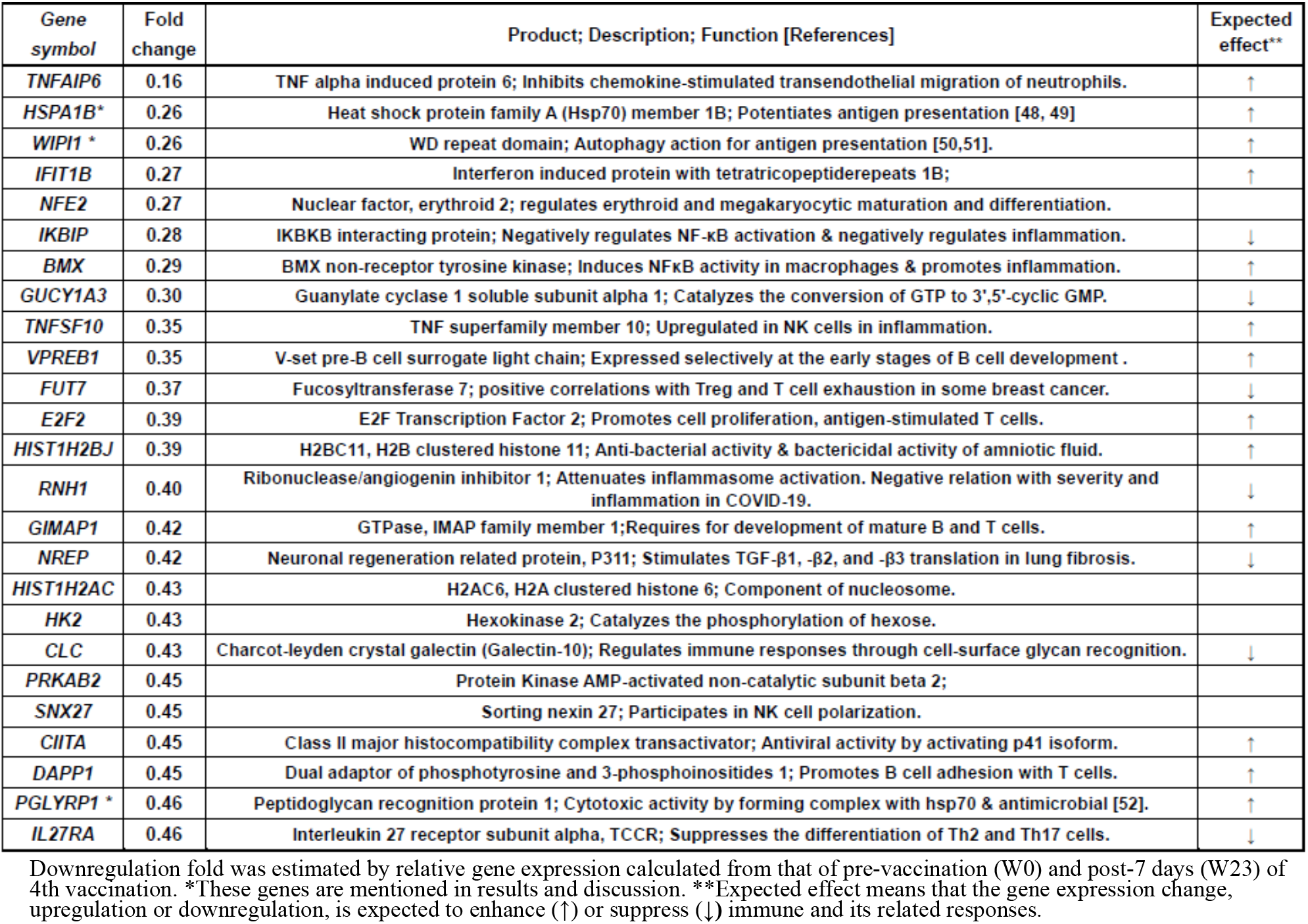
Immune-related genes downregulated less than 0.5-fold in Poly(I:C) group alone.

**Table 6.**
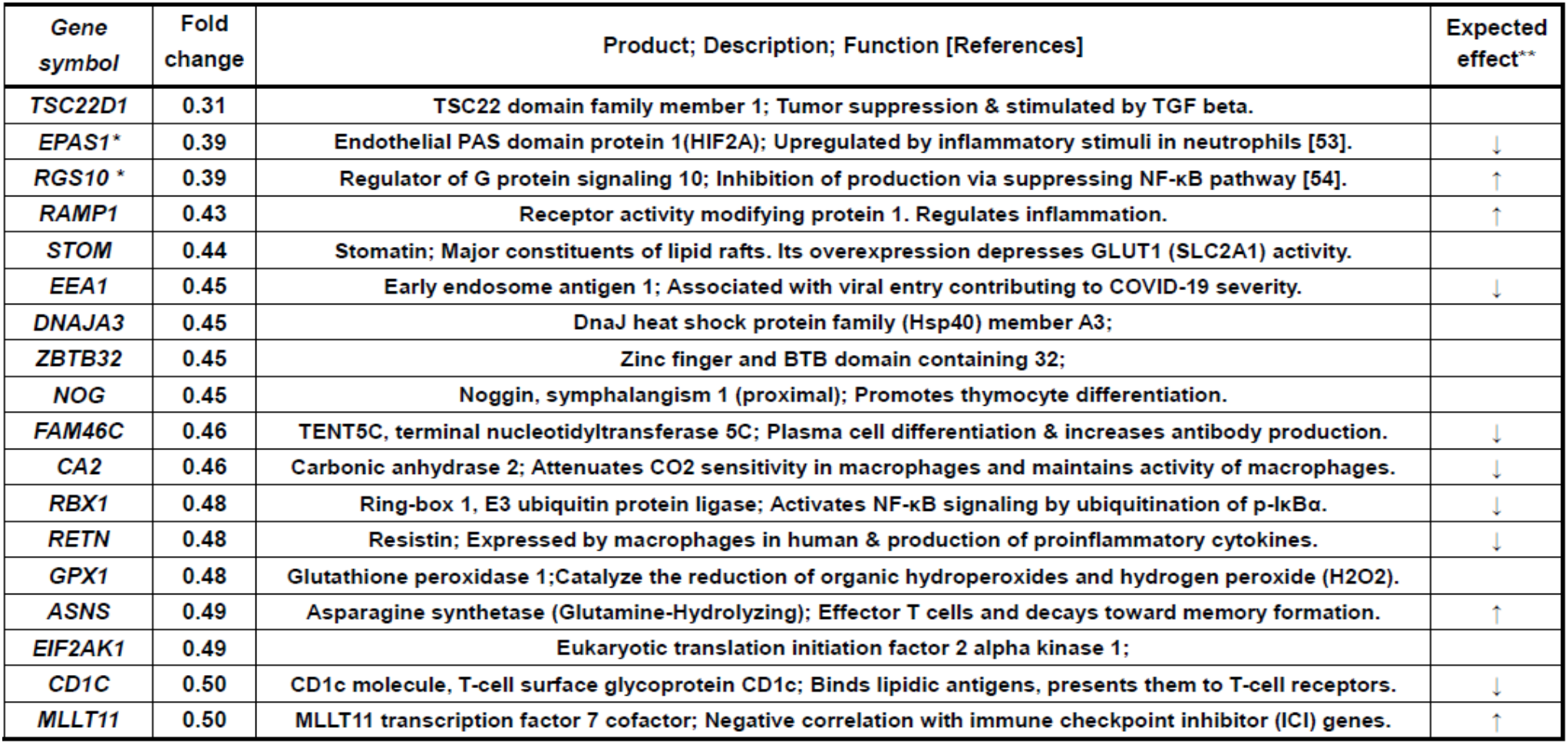
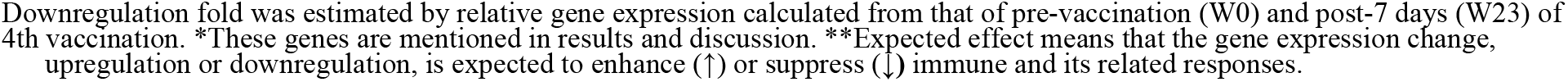
Immune-related genes downregulated less than 0.5-fold in AddaS03 group alone.

| Gene symbol | Fold change | Product; Description; Function [References] | Expected effect** |
| --- | --- | --- | --- |
| *TSC22D1* | 0.31 | TSC22 domain family member 1; Tumor suppression & stimulated by TGF beta. |  |
| *EPAS1\** | 0.39 | Endothelial PAS domain protein 1(HIF2A); Upregulated by inflammatory stimuli in neutrophils [53]. | ↓ |
| *RGS10\** | 0.39 | Regulator of G protein signaling 10; Inhibition of production via suppressing NF-κB pathway [54]. | ↑ |
| *RAMP1* | 0.43 | Receptor activity modifying protein 1. Regulates inflammation. | ↑ |
| *STOM* | 0.44 | Stomatin; Major constituents of lipid rafts. Its overexpression depresses GLUT1 (SLC2A1) activity. |  |
| *EEA1* | 0.45 | Early endosome antigen 1; Associated with viral entry contributing to COVID-19 severity. | ↓ |
| *DNAJA3* | 0.45 | DnaJ heat shock protein family (Hsp40) member A3; |  |
| *ZBTB32* | 0.45 | Zinc finger and BTB domain containing 32; |  |
| *NOG* | 0.45 | Noggin, symphalangism 1 (proximal); Promotes thymocyte differentiation. |  |
| *FAM46C* | 0.46 | TENT5C, terminal nucleotidyltransferase 5C; Plasma cell differentiation & increases antibody production. | ↓ |
| *CA2* | 0.46 | Carbonic anhydrase 2; Attenuates CO2 sensitivity in macrophages and maintains activity of macrophages. | ↓ |
| *RBX1* | 0.48 | Ring-box 1, E3 ubiquitin protein ligase; Activates NF-κB signaling by ubiquitination of p-IκBα. | ↓ |
| *RETN* | 0.48 | Resistin; Expressed by macrophages in human & production of proinflammatory cytokines. | ↓ |
| *GPX1* | 0.48 | Glutathione peroxidase 1; Catalyze the reduction of organic hydroperoxides and hydrogen peroxide (H2O2). |  |
| *ASNS* | 0.49 | Asparagine synthetase (Glutamine-Hydrolyzing); Effector T cells and decays toward memory formation. | ↑ |
| *EIF2AK1* | 0.49 | Eukaryotic translation initiation factor 2 alpha kinase 1; |  |
| *CD1C* | 0.50 | CD1c molecule, T-cell surface glycoprotein CD1c; Binds lipidic antigens, presents them to T-cell receptors. | ↓ |
| *MLLT11* | 0.50 | MLLT11 transcription factor 7 cofactor; Negative correlation with immune checkpoint inhibitor (ICI) genes. | ↑ |

Downregulation fold was estimated by relative gene expression calculated from that of pre-vaccination (W0) and post-7 days (W23) of 4th vaccination. \*These genes are mentioned in results and discussion. \*\*Expected effect means that the gene expression change, upregulation or downregulation, is expected to enhance (↑) or suppress (↓) immune and its related responses.

These included six marked genes; ***TSC22D1***: TGF-beta pathway activation, ***EPAS1***: stimulation of innate immunity via hypoxia response, ***RGS10***: NF-kB pathway inhibition, ***ZBTB32***: repression of recall of memory B-cell, ***RBX1***: activation of NF-kB pathway, and ***CD1C***: antigen presentation to T-cells.

This up- or downregulated gene expression could link to the enhancement or suppression of innate and acquired immune responses elicited by a sublingual vaccine formulated with RBD antigens and the Poly(I:C) or AddaS03 adjuvant.

## 4. Discussion

An adjuvant is critical for the safety and efficacy of a mucosal vaccine. Poly(I:C) is dsRNA, a TLR3 ligand that activates immune and proinflammatory responses [55]. The dsRNA mimics viral infection through binding endosomal TLR3 and cytosolic receptors, the retinoic acid-inducible gene I, and melanoma differentiation-associated gene 5 [56]. Poly(I:C) is a potent vaccine adjuvant that activates antigen-presenting cells, particularly dendritic cells [57, 58]. Poly(I:C)-mediated TLR-3 activation leads to proinflammatory cytokine, IFN, IL-15, and NK cell production [59]. However, clinical use of Poly(I:C) as a vaccine adjuvant has been unapproved except for limited use in cancer. Nonetheless, studies to develop its use as a vaccine adjuvant are progressing in preclinical and clinical fields [60]. The Poly(I:C)-mediated proinflammatory cytokines production and related factors were primarily reported in studies using nasal vaccination in mice [61, 62].

In the present study using non-human primate model, the Poly(I:C) adjuvant for sublingual vaccine use provided the same safety results as those of AddaS03 when examining complete blood count, biochemical blood test, and plasma CRP. The Poly(I:C) adjuvant is also considered safe compared to gene expression analyses with those of AddaS03. The Poly(I:C) adjuvant displayed the same gene expression profiles as those of AddaS03 in proinflammatory cytokines and its related factors: IL12a, IL12b, GZMB, IFN-alpha1, IFN-beta1, and CD69. Characteristically the gene expression of these cytokines returned to the pre-vaccination level at seven days post-vaccination, as did AddaS03, suggesting the safety evidence of the Poly(I:C) adjuvant. In previous research, the Poly(I:C)-mediated proinflammatory cytokines and related factors production were primarily reported in studies using nasal vaccination in mice [61, 62]. Differences in the immune system between rodents and primates were remarked by genome-based evidence [63]. Poly(I:C) is the most effective type I IFN inducer among TLR agonists [64]; however, its marked proinflammatory cytokine pathway activation may be overestimated. Moreover, the Poly(I:C)-mediated adverse events differed at nasal and sublingual sites. Previous studies on Poly(I:C)-mediated vice-reactivity are insufficient to establish the administration route via the nasal site and use of the rodent model.

Sublingual vaccination formulated with Poly(I:C) or AddaS03 adjuvant produced RBD-antigen specific IgA antibodies in the pituita and saliva and specific IgA and IgG antibodies in the blood. These indicate that the sublingual vaccine activated the local mucosal and systemic immune systems. The different specific antibody production patterns at the third vaccination were observed between saliva/nasal washings and plasma. These findings suggest that distinct immune responses were evoked by sublingual vaccination at local mucosal and systemic sites. The SARS-CoV-2 neutralizing antibody was detected in plasma samples from the Poly(I:C) and AddaS03 groups. However, no neutralizing antibody activity was observed in the saliva or nasal washings from both groups. This negative result could be caused by the SARS-CoV-2 anti-RBD Antibody Profiling kit used. The kit is based on the binding event between SARS-CoV-2 RBD and the angiotensin-converting enzyme 2 (ACE2) protein on the host cell surface (press release by Keio University on September 30, 2020). The soluble form of ACE2 exists in plasma, saliva and pituita [65, 66] and could inhibit the binding of SARS-CoV-2 RBD and ACE2 in the kit; hence the low activity of plasma and false-negative saliva and nasal washings occurred.

Sublingual vaccination formulated with Poly(I:C) or AddaS03 adjuvant had the same effect on Th1, Th2 induction, or both, judging from INF-alpha, IFN-gamma, and IL-12 production. The Poly(I:C)-formulated sublingual vaccine was more potent than AddaS03 for Th17 induction because of its 1.5-fold higher IL-17 production than AddaS03. Immune responses eliciting the Th17 subset play a critical role in vaccine-mediated protection against pathogens. This Th17 response, accompanied by neutralizing IgA antibody production, was reported in the sublingual influenza vaccine [67]. Excess Th17/IL17 response causes inflammation by activating the IL-17 receptor-mediated Act1/TRAF6 pathway [68]. Thus, moderate Th17 induction observed in the present study is the preferable mechanism of action of the sublingual vaccine formulated with Poly(I:C) adjuvant.

DNA microarray analyses are a high throughput method to investigate molecular mechanisms on the gene expression level. In the present study, we employed the DNA microarray technique to elucidate molecular events in the immune system and related responses mediated by sublingual vaccination formulated with the Poly(I:C) or AddaS03 adjuvant. The DNA microarray was performed using WBC RNA because WBCs are available in clinical studies to compare with current preclinical studies performed on non-human primates. A sublingual vaccine formulated with Poly(I:C) or AddaS03 adjuvant induces gene up- or downregulation of genes associated with immunity and related responses in the innate and acquired immune systems. These up- or downregulated gene expressions is expected to affect the enhancement or suppression of the immune responses.

In the point of view of enhancing effect to immune responses, *CCL7, CCL2, CXCR4, PFKFB3, JUN, KLHL2, PTX3, FADD*, and *ETV6* were upregulated by Poly(I:C) and AddaS03 adjuvanted vaccines (Table1). Coding products of these genes have an accelerating effect on the immune and its related responses; CCL7 and CCL2: chemokines to participate in monocyte migration to inflammatory sites from the bone marrow [17], CXCR4: chemokine receptor having a crucial role in B-cell generation [22], PFKFB3: a key enzyme to metabolize glucose and is associated with T-cell activation, macrophage survival, and neutrophil activation [23-25], Jun: AP-1 formation resulting in both T-cell activation and helper T-cell differentiation and stimulation of the CCR2 gene expression [26, 27], KLHL2: transcriptional factor in inducing *JUN* gene expression [31], PTX3: IL-6-induced acute inflammatory response protein [28], FADD: adaptor for apoptosis; NF-kB activation; and T-cell activation [29], and ETV6: stimulation of antigen memory in memory T-cells [30]. *C15orf48* and *SLC2A1* were upregulated by the Poly(I:C) adjuvanted vaccine (Table 2). These codings relate to the following immune reactions; C15orf48: stimulates the inflammatory reaction by miR-147b derived from C15orf48 mRNA [34, 35], and SLC2A1 (GLUT1): stimulation of glucose intake into T-cells for its activation [20]. The AddaS03 adjuvanted vaccine induced upregulated gene expression of *IFIT1, ITIF2, ITIF3, GBP1*, and *ISG15T* (Table 3). Their codings are associated with the following functions; IFITs: anti-virus action through inhibiting cap-dependent translation by binding to eITF3 [40], GBP1: Innate immune response to tuberculosis [41], and ISG15: Cytokine production for NK activation [42]. Concerning the enhancing effects on immune responses, we observed downregulated gene expression by vaccinations formulated with Poly(I:C) or AddaS03 adjuvant (Tables 4∼6). These genes were *ITGB5, PGLYRP1*, and *RGS10* for Poly(I:C) or AddaS03 adjuvanted, Poly(I:C) adjuvanted, and AddaS03 adjuvanted, respectively. Their codings were reported to have the following regulatory roles; ITGB5: macrophage polarization to antiinflammatory M2 acceleration [44], PGLYRP1: cytotoxic activity to target cells by forming a complex with hsp70 [52], and RGS10: inhibiting inflammation-mediated cytokine production via suppressing NF-κB pathway [54].

In the viewpoint of suppressive effect to immune response, *TNFRSF12A, RGS1*, and *SLA* were upregulated by both of Poly(I:C) and AddaS03 adjuvanted vaccines (Table1). The coding products of these genes are expected to have suppressive effect to on the immune and its related responses; TNFRSF12A: inhibition of T-cell proliferation and activation [18], RGS1: T-cell exhaustion [19, 20], and SLA: inhibits T-cell receptor-mediated signal transduction [21]. The Poly(I:C) adjuvanted vaccine also mediated upregulated gene expression of *ZNF165, EDN1, PRDM1, NR4A*, and *EWSR*1(Table 2). Their coding products have the following functions; ZNF165 and EDN1: repressing B-cell function by activating the TGF-beta pathway [32, 33], PRDM1: Treg differentiation accompanying inducing *FOXP3* expression [36], NR4A3: pivotal role in T-cell exhaustion [37], and EWSR1: inhibiting B-cell response for antigen-specific plasma cell generation [38]. The AddaS03 vaccine induced upregulated gene expression of *PDK4* and *GAB2* (Table3), whose codings are related to the following actions; PDK4: B-cell differentiation inhibition through CD40 production and cell surface expression repression [39] and GAB2: inhibition of T-cell activation by reducing IL-2 production [43]. Concerning the suppressive effects on immune responses, we also found downregulated gene expression of *APQ1, APQ3*, GP9, and *GP1BB* by the Poly(I:C) and AddaS03 adjuvanted vaccines (Table 4). Their codings have the following immune-stimulating functions; ApQ1: IL-beta production in macrophages and infiltration of neutrophils [47], APQ3: IL-6 and TNF-alpha production in macrophages [45], and GP9 and GP1BB: formation of the neutrophil extracellular trap through activation of the blood clotting system [46]. The suppressive effect was observed by downregulated gene expression mediated with the Poly(I:C) or AddaS03 adjuvanted vaccine (Table 5, 6). These downregulated genes were *HSPA1B, WIPI1*, and *EPAS1*; HSPA1B: potentiation of antigen presentation [48, 49], WIPI1: autophagy action for antigen presentation [50, 50], and EPS1: induction of hypoxia response of immune cells [53].

Therefore, a sublingual vaccine formulated with Poly(I:C) or AddaS03 adjuvant affected transcriptional regulation to up- or down-regulate many genes related to enhancing or suppressing the immune responses. Thus, the sublingual vaccination appeared to yield a balance between enhancing and suppressing the immune responses by regulating their gene expression. Sublingual vaccination with the Poly(I:C) or AddaS03 adjuvant probably has some counterbalance like a “Yin and Yang” manner to brake and accelerate the immune responses. Furthermore, the sublingual vaccinations lead to unique suppressive effects such as T-cell proliferation inhibition (*TNFRSF12A*), T-cell exhaustion (*RGS1* and *NR4A*), Treg differentiation (*PRDM1*), B-cell function repression (*EDN1*), and B-cell response inhibition (*EWSR1*). The previous paper reported that sublingual-induced oral tolerance generated Treg generation [69]. Sublingual administration of Ag/CTB conjugates induced immunological tolerance to generate Treg [70]. Induction of tolerance via the sublingual route was also seen in the review paper [71]. Therefore, as mucosal immunization, including sublingual vaccine administration, result in some immune suppression: tolerance ; Treg generation ; -and T-cell exhaustion, the sublingual route is selected for allergy immunotherapy. Furthermore, sublingual vaccination formulated with the Poly(I:C) adjuvant elicited atypical immune suppression. T-cell exhaustion accompanied by increased co-suppressive receptor PD-1 production declined cell proliferation. However, the sublingual vaccination induced *RGS1* and *NR4A* upregulation but not *PCDI*, suggesting insufficient inhibition of T-cell proliferation via increased PD-1. Treg differentiation accompanied the *FOXP3* upregulation to promote the differentiation of naïve T-cells to Treg [36]. In contrast, this sublingual vaccination induced upregulated *PRDM1* expression but not *POXP3* nor *KLF2*, indicating incomplete RPDM1-promoted T-cell hyporesponsiveness. This unusual transcriptional regulation appears to be characteristic of Poly(I:C)-adjuvanted sublingual vaccination. This study is the first to mention atypical transcriptional regulation for immune tolerance; T-cell exhaustion; Treg differentiation; or T-cell hyporesponsiveness elicited by sublingual vaccination formulated with the Poly(I:C) adjuvant.

## 5. Conclusions

A sublingual vaccine formulated with RBD antigen and Poly(I:C) adjuvant had the same safety as the AddaS03 adjuvant. Furthermore, RT-qPCR of 6 inflammatory cytokines and ELISA of 4 cytokines indicated the safety of Poly(I:C) adjuvanted sublingual vaccine. The Poly(I:C) and AddaS03 adjuvanted vaccine evoked RBD-specific IgA antibody production in nasal washings, saliva, and plasma. SARS-CoV-2 neutralizing antibody was detected in plasma, suggesting that both adjuvanted vaccines had the immune potential to protect against SARS-CoV-2. A sublingual vaccine formulated with Poly(I:C) or AddaS03 adjuvant elicited unique transcriptional regulation in a “Yin and Yong” manner. The transcriptional regulation induced by sublingual vaccination could enhance and suppress immune responses via up- or downregulation of their genes. The Poly(I:C) adjuvanted sublingual vaccination especially induced atypical up- or downregulated gene expression associated with immune suppression/tolerance, resulting in incomplete Treg differentiation and T-cell exhaustion.

## Author Contributions

Conceptualization, T. Y. and S.N.; methodology, F.M.; investigation, F.M and S.N.; resources, T. Y. and M.T.; data curation, F.M.; writing-original draft preparation, S.N. and F.M.; writing-review and editing, S.N. and T. Y.; visualization, F.M.; preparation and arrange of project, K.W., A.K., and K.T.; supervision and project administration, T. Y. and M.T.; funding, T. Y.; All authors have read and agreed to the published version of the manuscript.

## Institutional Review Board Statement

This study was conducted according to the guidelines of Institutional Animal Care and Committee Guide of Intelligence and Technology Lab, Inc. (ITL) based on the Guidelines for Proper Conduct of Animal Experiments and approved by the Animal Care Committee of the ITL (approved number: AE2021001, data: 7 July 2021). This study was also approved by the ITL Biosafety Committee (approved number: BS202100, date: 7 July 2021).

## Informed Consent Statement

Not applicable.

## Data Availability Statement

Data are available from S.N. upon reasonable request.

## Acknowledgments

We thank to Dr Makoto Hirano for his excellent informatics works and Kazuhiro Kawai for his invaluable technical assistance.

